# Striatal ups or downs? Neural correlates of monetary reward anticipation, cue reactivity and their interaction in alcohol use disorder and gambling disorder

**DOI:** 10.1101/2022.09.05.506605

**Authors:** Tim van Timmeren, Ruth J. van Holst, Anna E. Goudriaan

## Abstract

Striatal dysfunction is a key characteristic of addictive disorders, but neuroimaging studies have reported conflicting findings. An integrative model of addiction points to the presence or absence of addiction-related cues as an explanation for striatal hypo-or hyperactivations, respectively, but has never been directly tested. Here, we developed a novel paradigm to investigate striatal activation during monetary reward anticipation in the presence versus absence of addiction-related pictures using functional MRI. Across two studies, we compared 46 alcohol use disorder (AUD) patients with 30 matched healthy controls; and 24 gambling disorder (GD) patients with 22 matched healthy controls. During monetary reward anticipation, hypoactivation of the reward system was seen in AUD individuals compared to HCs. Additionally, a behavioral interaction was seen where gambling cues made participants, across groups, respond faster for bigger, but slower for smaller rewards. However, no striatal differences were seen between the participants with AUD or GD and their matched controls. In sum, these findings suggest that striatal dysfunction is a key but heterogeneous mechanism within both AUD and GD and indicates an important but complex role for addiction-related cues in explaining striatal dysfunction in addiction.

## INTRODUCTION

Patients with addictive disorders often show disrupted striatal reward processing (Bjork, Smith, Chen, & Hommer, 2012; Blum et al., 2000; Volkow & Morales, 2015). Findings have been inconsistent however, as both hypo- and hyperactivations have been reported (Clark, Boileau, & Zack, 2019; Leyton & Vezina, 2013; Limbrick-Oldfield, Van Holst, & Clark, 2013). Previous findings have been interpreted in the context of several addiction-theories, with largely incompatible predictions about the direction of the striatal disruption. A *hypo*active reward system (and related anhedonia) has been described either as a predisposition for the development of addictive behaviors (Blum et al., 2000) or as a consequence of chronic drug use and receptor down-regulation (Goldstein & Volkow, 2011; Koob & Le Moal, 2008), possibly together with the recruitment of ‘anti-reward’ systems (Koob & Le Moal, 2005). Alternatively, a *hyper*active reward system has been described either as a vulnerability factor reflecting increased sensitivity to high rewards driving impulsive behaviors (Bjork et al., 2012) or as a result of incentive sensitization for environmental stimuli that become conditioned with the rewarding effects of the addictive behavior (Robinson & Berridge, 2008).

Over the past decades, each of these theories has found scientific support, resulting in ample but inconsistent evidence for dysfunctions in the human reward system in addicted populations. It has been proposed that these seemingly contradictory findings may be integrated by considering the *presence* versus *absence* of addiction-related cues (Leyton & Vezina, 2013, 2014). Stimuli regularly associated with the addictive behavior become conditioned through repeated association with their rewarding effects – ultimately leading to sensitized neurobiological responses and craving (Vezina & Leyton, 2009). Hyperactive striatal motivational states thus develop in the presence of addiction-related cues, a phenomenon known as cue-reactivity. Simultaneously, a progressively diminished interest towards rewards unrelated to the addiction is reflected by a hypoactive reward system. Indeed, a review of the human substance use and gambling disorder literature suggests that many inconsistencies in the literature can be explained by factoring in addiction-related cues (Leyton & Vezina, 2013).

However, the presence of both striatal ‘ups’ and ‘downs’ within addicted patients has never been directly tested in the context of a single experimental paradigm. Such a study would not only be able to directly investigate the pervasiveness of ‘striatal ups and downs’ in addicted populations, but would also allow for an evaluation of the interaction between addiction-related cues (Freeman, Morgan, Beesley, & Curran, 2012) and monetary reward anticipation. The latter is often studied using the Monetary Incentive Delay Task [MIDT] (Balodis & Potenza, 2015; Beck et al., 2009; Knutson, Westdorp, Kaiser, & Hommer, 2000; Luijten, Schellekens, Kühn, Machielse, & Sescousse, 2017; Wrase et al., 2007). Cue-reactivity paradigms, during which participants are presented with addiction-relevant, have been frequently used in both substance (Zilberman, Lavidor, Yadid, & Rassovsky, 2019) and behavioral addictions (Starcke, Antons, Trotzke, & Brand, 2018). Addiction-related cues are known to increase motivation and modify readiness to respond for substances (Leyton & Vezina, 2013), but whether this effect is specific to drug seeking or generalizes to responding for natural rewards is still an open question. For example, gambling cues may augment the anticipation of a monetary reward in GD patients, such that the presence of such cues increases performance, motivation and striatal activity for monetary rewards (van Holst, van der Meer, et al., 2012; van Holst, Veltman, Van Den Brink, & Goudriaan, 2012). Such an effect might be indicative of (general) Pavlovian-to-Instrumental Transfer (PIT), a mechanism thought to be central to addiction (Everitt & Robbins, 2005) by which cues linked to some other reward could motivate instrumental behavior to an unrelated reward. Moreover, it is unclear if striatal ‘ups’ in response to addiction-cues correlate with striatal ‘downs’ during the processing of natural rewards within addicted individuals: they may be dependent factors that simultaneously develop with addiction (Volkow, Koob, & McLellan, 2016), or independent (risk-)factors constituting different addiction-subtypes.

Here we assessed monetary reward anticipation in the presence and absence of addiction-related stimuli to directly address these open questions. We adapted the widely used MIDT (Knutson et al., 2000) to include neutral and addiction-related cues during the anticipation phase and adopted a full-factorial design with the factors: monetary reward-type (low and high) and cue-type (neutral and addiction-related). This enabled us to separately study (i) general (monetary) reward anticipation, (ii) processing of addiction-related cues and (iii) their interaction, in both addicted and healthy control groups. We conducted two studies: one in patients with alcohol use disorder [AUD] and one in patients with gambling disorder [GD]. Both groups were separately matched to healthy control participants [HCs] and tested during fMRI-scanning to assess striatal functioning during the MIDT task in the presence or absence of addiction-related pictures. We hypothesized patient groups to show blunted striatal activity during monetary reward anticipation (i.e. striatal ‘downs’) in the absence of addiction-related cues, but increased motivation and neural reward-processing (i.e. striatal ‘ups’) in the presence of addiction-related cues compared to HCs.

## METHODS & MATERIALS

More details about the methods and results are provided in the Supplementary Material.

### Participants

All patients received treatment and were recruited from a local addiction treatment centre. AUD patients were detoxified (>2 weeks) and recently diagnosed with AUD without Axis 1 comorbidity. GD patients were included if they were recently diagnosed with and started therapy for GD but were not obliged to abstain from gambling. Patient groups were recruited through advertisements and our subject-database. All subjects underwent the MINI structured psychiatric interview (Sheehan, Lecrubier, & Sheehan, 1998), to confirm the absence of psychiatric disorders (except for DSM-5 AUD/GD in the AUD/GD patient group, respectively). Participants were included after meeting the inclusion criteria (see Supplementary Methods) and were reimbursed with 50 euros plus additional task earning (∼20 euros).

### Experimental procedure

After providing written consent, participants underwent ∼1 hour of interviews, questionnaires and cognitive tests. These data were collected as part of a more extensive study protocol which included another fMRI task and a resting-state fMRI scan (total scanning duration was 90 minutes), data of which have been (van Timmeren et al., 2020; van Timmeren, Zhutovsky, van Holst, & Goudriaan, 2018) or will be presented elsewhere. All fMRI sessions commenced in the afternoon (between 12:15 and 5:30 pm).

### Experimental paradigm

To investigate the effects of reward anticipation, cue-reactivity and their interaction, we adapted the MIDT (Knutson, Adams, Fong, & Hommer, 2001) to include addiction-related cues (alcohol- and gambling-relevant cues in the AUD-/GD-study, respectively). A total of 28 alcohol and neutral pictures were selected from The Geneva Appetitive Alcohol Pictures database (Billieux et al., 2011) supplemented by pictures from a previous study (Sjoerds, van den Brink, Beekman, Penninx, & Veltman, 2014) and pictures from the internet. The 28 gambling pictures were selected from a previous study (Goudriaan, de Ruiter, van den Brink, Oosterlaan, & Veltman, 2010) from the internet. Neutral pictures were matched (independently to the alcohol and gambling pictures) for setting, color-distribution, and complexity.

We used a 2×2 full-factorial design with reward magnitude (Big reward=50cent coin and Small reward=1cent coin) and cue-type (addiction-related and neutral background pictures) as factors, resulting in four conditions: ‘Big reward/Addiction cue’ [BA], ‘Big reward/Neutral cue’ [BN], ‘Small reward/Addiction cue’ [SA] ‘Small reward/Neutral cue’ [SN]. The primary dependent measure was fMRI BOLD response during the MIDT reward anticipation stage. The task comprised 28 trials per condition; trials were presented pseudo randomly and the total task duration was ∼23 minutes. Figure 1a shows the experimental design, more detailed information about the procedure is included in the Supplementary information.

**Figure 1.**
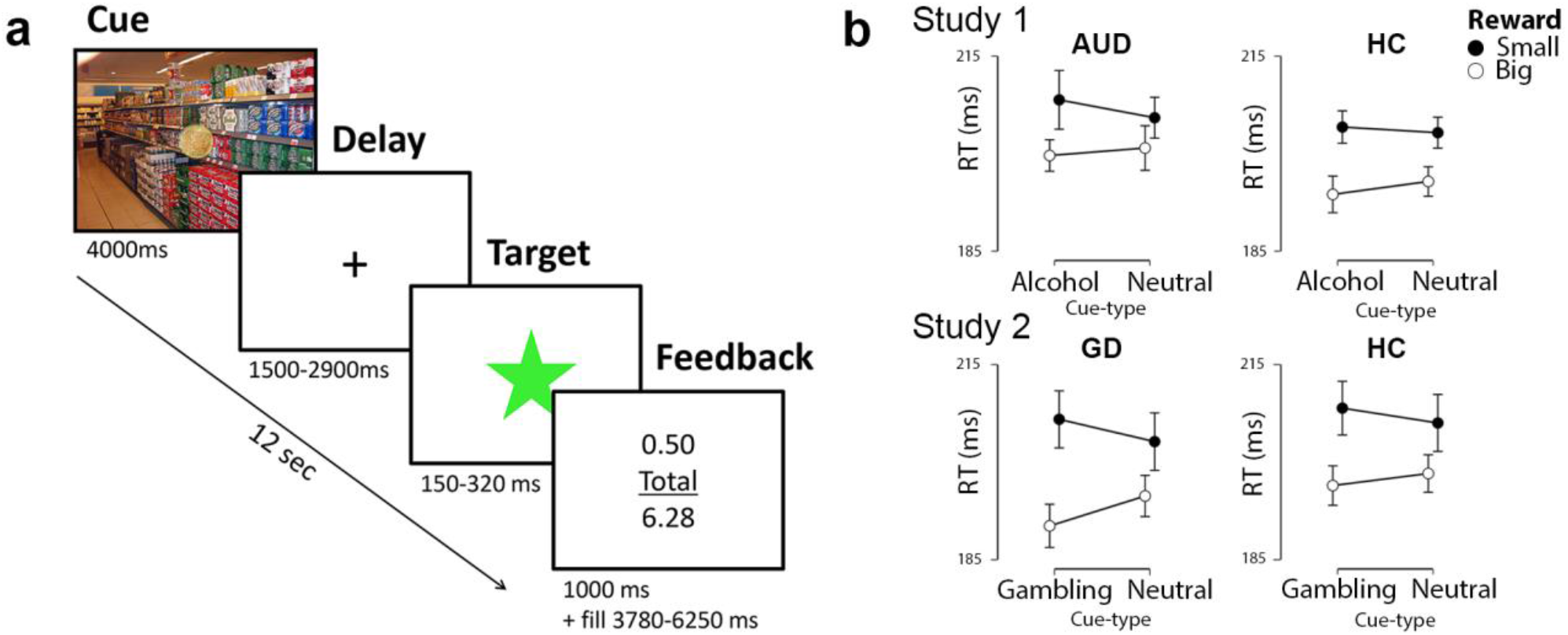
Schematic overview of the experimental design and behavioral (reaction time) results. (a) Participants were instructed to respond to a target as quickly as possible in order to gain monetary rewards. During each trial, participants could earn 1 or 50 cents, indicated by a coin overlaid on an addiction-related or neutral background picture (‘cue’). Next, a crosshair was shown for a variable period (‘Delay’) and participants were instructed to respond as fast as possible to the target. Feedback about current and cumulative earnings was provided, followed by a fixation cross before a new trial started. (b) Mean reaction time (in ms) on the task for each condition, showing faster responses for big than small rewards in both studies. Moreover, in the GD study, a significant interaction was found between reward- and cue-type, such that responses were faster for bigger rewards and slower for smaller rewards in the presence of gambling cues, whereas this relation was reversed in small reward condition. Error bars indicate 95% confidence interval.

Following fMRI acquisition, participants were asked to indicate how strongly each background-picture induced craving on a 7-point Likert scale (i.e. subjective craving). Technical failures resulted in missing craving data of six participants in the AUD study (three AUD patients and three HCs) and seven participants in the GD study (two GD patients and five HCs).

### Magnetic Resonance Imaging

#### Acquisition

Participants entered the 3T Phillips MRI-scanner in head-first supine position and were able to view the screen using a mirror attached to the head-coil. We acquired 595 T2*-weighted multiecho planar functional MRI volumes (voxel-size: 3mm^3^) for analysis. Additionally, a structural T1-weighted image (voxel-size: 1mm^3^) was collected. See Supplementary Materials for details.

#### fMRI analysis

All functional MRI data were analysed using SPM12 (Wellcome Trust Centre for Neuroimaging, London, United Kingdom). Raw multiecho fMRI data were first combined into single volumes using the PAID-method (Poser, Versluis, Hoogduin, & Norris, 2006). Preprocessing of the fMRI data was identical to (van Timmeren et al., 2018) and involved motion correction, slice-time correction, co-registration, normalization and smoothing (see Supplementary Materials for details). A first-level general linear model was constructed for each participant, including individual regressors for all four conditions (4 second box-car function at stimulus-onset). The feedback phase was not considered here because there was no interaction with addiction-related cues; outcome (win/loss) and key presses were included as regressors of no interest. Realignment parameters were entered as six nuisance regressors and low frequency drifts were removed using a high-pass filter (128-s cutoff). Regressors were convolved with a canonical hemodynamic response function.

Three first-level contrast images were constructed for each participant to assess the effect of (1) reward type (monetary reward anticipation = big versus small reward: [BA+BN]>[SA+SN]); cue type (cue-reactivity = alcohol versus neutral cues: [BA+SA]>[BN+SN]); and (3) their interaction ([BA+SN]>[BN+SA]). Three *a priori* striatal regions of interests were derived from the Oxford-Imanova Striatal Structural Atlas (http://fsl.fmrib.ox.ac.uk): bilateral ventral striatum, caudate and putamen. For each participant, parameter estimates were extracted and averaged across voxels for the three ROIs separately to investigate regional activation for the relevant contrasts (i.e., monetary reward anticipation, cue-reactivity and their interaction effect). Additionally, single-subject contrast images were entered into second-level random-effects analysis, comparing within-group activation (one-sample *t* tests) and between-group differences (two-sample *t* tests). These whole brain analyses were additionally conducted to describe any non-striatal group differences for the monetary reward anticipation, cue-reactivity and their interaction. These results were corrected for whole-brain familywise error [FWE] (α<.05, voxelwise p<.001) and anatomical brain regions were identified using the Automated Anatomical Labelling [AAL] atlas in SPM12 (Tzourio-Mazoyer et al., 2002).

### Exploratory analyses

#### Addiction severity, chronicity and abstinence

We additionally explored whether individual differences in striatal functioning in the clinical groups were associated with several clinical measures found to be relevant in previous studies: in both groups, we looked at craving levels, which have been related to increased ventral striatal cue-reactivity (Filbey et al., 2008) and blunted ventral striatal monetary reward processing (Wrase et al., 2007) in AUD, while in GD craving levels have been found to correlate with increased insular cue-reactivity (Goudriaan et al., 2010; Limbrick-Oldfield et al., 2017). We also looked at duration of abstinence, which has previously been associated with striatal cue-reactivity (although positively in AUD (Li et al., 2014), but negatively in GD (Limbrick-Oldfield et al., 2017)). Finally, we looked at addiction severity and duration, which have been related to striatal cue-reactivity (Claus, Ewing, Filbey, Sabbineni, & Hutchison, 2011; Ihssen, Cox, Wiggett, Fadardi, & Linden, 2011; Sjoerds et al., 2014; Vollstädt-Klein et al., 2010).

In the group of patients with AUD, the following outcomes were used: AUD severity (AUDIT, range 0-40; Saunders, Aasland, Babor, de la Fuente, & Grant, 1993), obsessive alcohol-related thoughts (OCDS, range 0-20; De Wildt et al., 2005) and lifetime alcohol intake (kg) (Skinner & Sheu, 1982). Within the GD group, GD severity was measured with the PGSI (Ferris & Wynne, 2001). In both groups, we looked at subjective craving (obtained by rating the pictures post-scanning, see Experimental procedure), duration of addiction problems (years) and abstinence (days). Pearson’s correlations were done between these factors and the three striatal ROIs during monetary reward anticipation and cue-reactivity. Considering the multitude of tests (involving 3 ROIs, 2 events of interest and a total of 9 clinical factors), we refrain from making any statistical inferences (i.e. using p-values) and only report results for moderate to high correlations (Pearson’s correlation coefficient > 0.3). All per test in the supplementary data.

#### Relation between striatal ‘ups’ and ‘downs’

To test whether increased striatal cue-reactivity (striatal ‘ups’) and diminished striatal monetary reward anticipation (striatal ‘downs’) were related to each other within individual patients, Pearson correlations were performed (for both patient groups separately) using the extracted parameter estimates for each of the three ROIs. Evidence for the null hypothesis (i.e. no relation between the two factors) were substantiated by Bayes Factors using Bayesian correlations.

### Statistical Analysis

Statistical analysis was carried out using JASP, version 0.8.6 (JASP Team, 2018). Demographics and clinical data were analyzed for group differences with two-sampled t-tests and Pearson’s chi-square tests for each study separately. Non-normally distributed data were analyzed using Mann-Whitney *U*-tests for group comparisons. Mixed ANOVAs were used to analyze mean reaction times and number of hits, using reward magnitude (big or small) and cue type (addiction or neutral) as within-subject factors and group (patients or controls) as between-subject factor. Striatal group differences were analyzed using independent t-tests by taking the extracted parameter estimates for the three main fMRI contrasts: reward anticipation, cue-reactivity and their interaction effect.

A significance threshold of p<0.05 (two-tailed) was considered significant. Striatal analyses were additionally conducted using corresponding Bayesian analyses, using JASP’s default Cauchy prior (0.707). The resulting Bayes Factor_10_ (BF_10_) indicates how much more likely the data are under the alternative hypothesis (H1) than under the null hypothesis (H0). We also report the BF_01_, which quantifies the relative evidence in favor of the null hypothesis, or in other words the amount of support for the *absence* of an effect. BF between 1 and three is considered to reflect anecdotal evidence, BF > 3 reflects substantial support and values larger than 10 reflect strong support (Wetzels et al., 2011). See Wagenmakers et al., 2018 for an introduction into Bayesian hypothesis testing.

#### Ethics

The study was approved by the local Ethical Review Board of the Academic Medical Center, University of Amsterdam, the Netherlands (2014_345). All subjects provided written informed consent.

## RESULTS

Results are reported for the AUD and GD study separately. All main neuroimaging results are available online at https://neurovault.org/collections/4199/.

## STUDY 1: AUD

### Demographics and clinical characteristics

The groups were matched on age (AUD: mean=46.5, SD=10.8; HCL mean=44.8, SD=9.9), handedness, gender, years of education and IQ. As expected, the AUD group had significantly more smokers (p<0.001) and scored higher on all factors related to alcohol use: AUDIT, lifetime alcohol intake, kg pure alcohol use and OCDS (all p<0.001). Demographics and clinical characteristics are summarized in Supplementary Table 1.

### Behavioral results

The repeated measures ANOVAs of reaction time and number of hits indicated a main effect of reward: as expected, participants were faster (F_1,74_=38.9; p<0.001, η^2^=0.34) and more accurate (F_1,74_=20.2; p<0.001, η^2^=0.21) for big compared to small rewards (Figure 1B and S1). No significant main or interaction effects of cue-type or group were found.

For the cue-induced craving ratings there was a group by cue-type interaction (F_1,68_=32.2, p<0.001). Post-hoc tests revealed this interaction was driven by significantly higher craving ratings for alcohol cues in AUD patients (mean=3.49, SD=1.92) compared to neutral cues in AUD patients (mean=1.44, SD=0.58; t_42_=7.40 p<0.001) and compared to alcohol cues in HCs (mean=1.49, SD=0.70; t_68_=5.18 p<0.001).

### Imaging results

Figure 4b includes a visual overview of average ventral striatal activity on the different task conditions for each group.

#### Reward anticipation

In line with our hypothesis, AUD patients showed significantly decreased activity, relative to HCs, while anticipating big versus small monetary rewards in all three ROIs: the ventral striatum, putamen and caudate (all p<0.001; see Table 1). Whole brain sensitivity analyses further showed pallidum, insula, hippocampus and supplementary motor area extending to the right middle and medial cingulate and inferior OFC (Figure 3A).

**Table 1.**
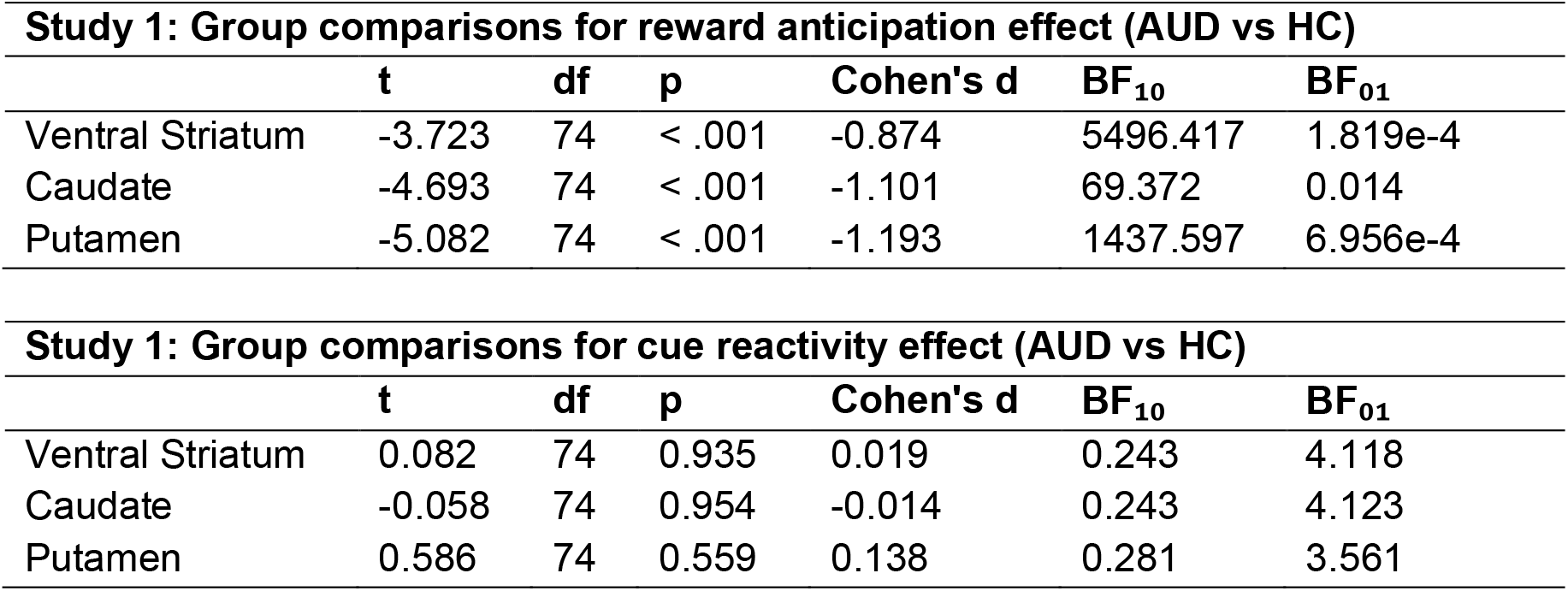
(Bayesian) independent samples t-tests were used to test group comparisons between AUD patients and HCs on the effect of reward anticipation (big > small) and cue reactivity (alcohol > neutral cues), for all three regions of interest. BF: Bayes Factor, see Statistical Analysis for more information.

#### Cue-reactivity

Contrary to our hypothesis, no significant group difference in striatal activation (see Table 1) or on the whole brain level was found for the contrast comparing alcohol with neutral cues. Within the AUD group alone, this cue-reactivity contrast revealed significantly increased activation in several clusters in the bilateral precuneus extending to the middle cingulate, the bilateral frontal superior medial extending to the anterior cingulate cortex [ACC], the left frontal inferior orbital cortex and several occipital regions (left angular and right lingual gyrus); see Figure 2B. In contrast, HCs showed increased activity only in the primary visual cortex. Thus, neural responses to alcohol cues in the AUD group were dissociable from neutral cues, but not significantly different from those in HCs.

**Figure 2:**
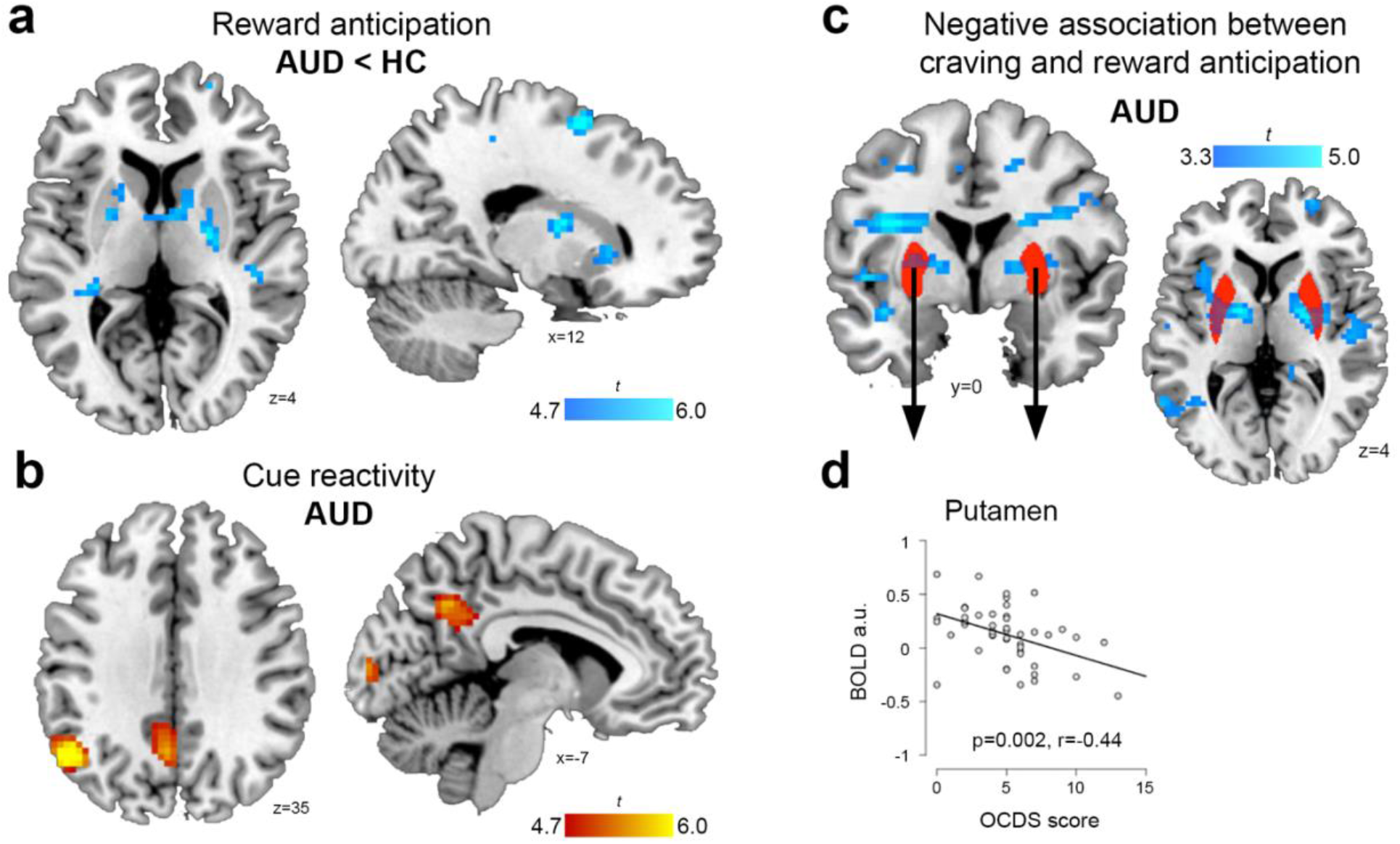
AUD patients showed decreased activity during monetary reward anticipation relative to HCs (a) and increased activity to addiction-related compared to neutral stimuli (b). A negative association was found between the level of obsessive alcohol-related thoughts (OCDS) and reward anticipation (big>small) in AUD (c&d). **(a)** Whole-brain statistical parametric map showing blunted monetary reward anticipation (big>small rewards) in AUD patients compared to HCs (AUD<HC). Results shown at p<0.05, FWE-corrected. **(b)** Cue-reactivity (alcohol>neutral cues) within AUD group. Results shown at p<0.05, FWE-corrected. **(c)** Whole-brain statistical parametric map for the correlation between OCDS scores and the big>small contrast, shown at p<0.001 uncorrected. **(d)** Correlation between OCDS and mean parameter estimates extracted from the putamen.

#### Reward x cue-type interaction

The reward-magnitude–by–cue-type interaction revealed no significant differences in BOLD activity between AUD patients and HCs, or within the group of AUD individuals, both within the striatal ROIs and on the whole-brain.

## Study 2: Gambling Disorder

The GD and HC groups were matched on age (GD: mean=35.5, SD=12.4; HC: mean=35.4, SD=15.8), gender, years of education, alcohol use and smoking status, but differed on handedness, IQ and factors related to gambling. Demographics and clinical characteristics are presented in Supplementary Table 2.

### Behavioral results

Similar to the results in Study 1, main effects of reward were seen for reaction time (F_1,44_=63.9; p<0.001, η^2^=0.59) and number of hits (F_1,44_=26.3; p<0.001, η^2^=0.37), driven by participants being faster and more accurate for big compared to small rewards (Figure 1B and S1). Moreover, significant reward*cue-type interactions were found for both reaction time (F_1,44_=4.7; p=0.035, η^2^=0.10) and hits (F_1,44_=4.6; p=0.037, η^2^=0.09): in the presence of gambling cues, GD and HC participants were faster when playing for big rewards, but slower when playing for small rewards. No significant main or interaction effects of group were found.

As expected, there was a group-by-cue-type interaction for the post-scan craving ratings (F_1,37_=12.6, p=0.001, *η*_p_^2^=.08), driven by higher ratings for gambling (mean=3.98, SD=2.65) compared to neutral cues (mean=1.32, SD=0.81;) in GD patients (t_21_=5.15, p<0.001) and compared to gambling cues in HCs (mean=1.75, SD=0.80; t_37_=3.34 p=0.002; neutral cues HCs: mean=1.24, SD=.05).

### Imaging results

#### Reward anticipation

Contrary to our hypothesis, no significant group difference in striatal activation was found in the contrast comparing anticipation of big monetary rewards versus small monetary rewards (Table 2). On the whole-brain level, GD patients showed decreased activation relative to HCs during reward anticipation in two clusters, with one peak in the left angular gyrus extending to the temporal middle and superior gyrus, and one peak in the right temporal superior gyrus extending to the temporal middle gyrus and hippocampus (Figure 3A).

**Table 2.**
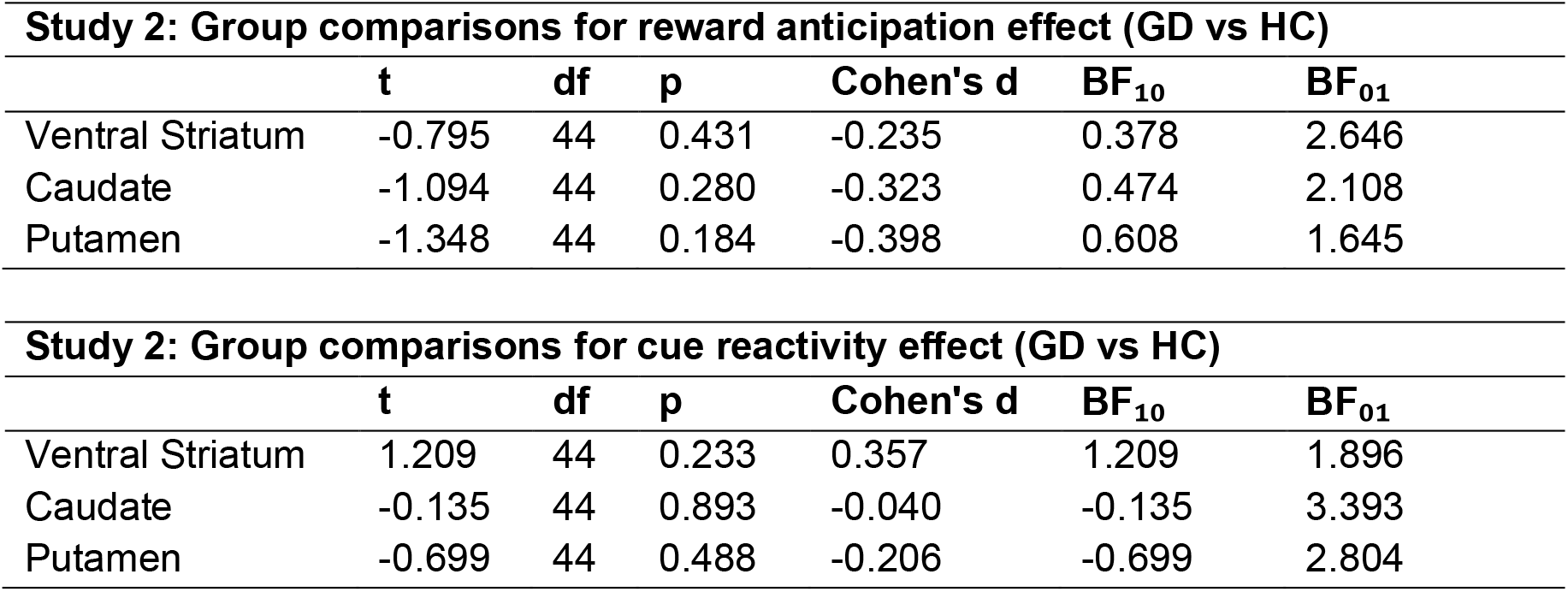
(Bayesian) independent samples t-tests were used to conduct group comparisons between GD patients and HCs on the effect of reward anticipation (big > small) and cue reactivity (gambling > neutral cues), for all three regions of interest. BF: Bayes Factor, see Statistical Analysis for more information.

| **Study 2: Group comparisons for reward anticipation effect (GD vs HC)** |  |  |  |  |  |  |
| --- | --- | --- | --- | --- | --- | --- |
|  | **t** | **df** | **p** | **Cohen's d** | **BF<sub>10</sub>** | **BF<sub>01</sub>** |
| Ventral Striatum | -0.795 | 44 | 0.431 | -0.235 | 0.378 | 2.646 |
| Caudate | -1.094 | 44 | 0.280 | -0.323 | 0.474 | 2.108 |
| Putamen | -1.348 | 44 | 0.184 | -0.398 | 0.608 | 1.645 |

| **Study 2: Group comparisons for cue reactivity effect (GD vs HC)** |  |  |  |  |  |  |
| --- | --- | --- | --- | --- | --- | --- |
|  | **t** | **df** | **p** | **Cohen's d** | **BF<sub>10</sub>** | **BF<sub>01</sub>** |
| Ventral Striatum | 1.209 | 44 | 0.233 | 0.357 | 1.209 | 1.896 |
| Caudate | -0.135 | 44 | 0.893 | -0.040 | -0.135 | 3.393 |
| Putamen | -0.699 | 44 | 0.488 | -0.206 | -0.699 | 2.804 |

**Figure 3.**
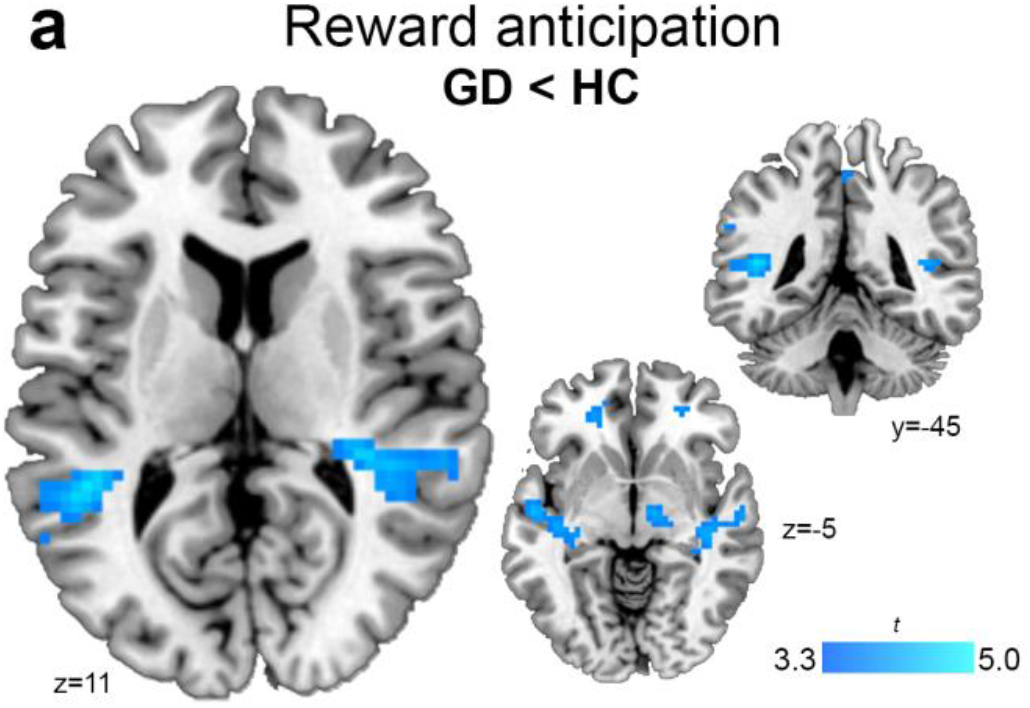
GD patients showed decreased activity during monetary reward anticipation. Whole-brain statistical parametric map showing blunted reward anticipation in gamblers compared to controls (GD<HC).

#### Cue-reactivity

Similar to the results from the AUD study, no significant differences were found between GD patients and HCs on the contrast comparing gambling with neutral cues, neither in the striatal ROIs (Table 2) nor whole-brain. Within the group of patients with GD, increased activation of the bilateral calcarine and lingual gyrus was seen during cue-reactivity (Figure S2).

#### Reward x cue-type Interaction

No significant differences were found for the reward-magnitude*cue-type interaction between the groups or within GD patients.

### Exploratory analyses

#### Relation striatal activity and craving, severity, chronicity, and abstinence

Tables with all correlation coefficients are included in the supplementary material (Suppl Tables 5-8). As we included those correlations entirely exploratory, it’s not meaningful to report p-values (Gelman & Loken, 2013), which is why we only present and interpret medium and larger effect sizes (r>0.3). A moderate (r=-0.44) negative correlation was seen between OCDS scores and reward anticipation in the putamen (Figure 2D). Thus, AUD patients who reported having more obsessive alcohol-related thoughts showed stronger hyporesponsive dorsal striatal activity during monetary reward processing. In the group of patients with GD, ventral striatal cue reactivity was moderately (r=0.44) correlated to the duration of gambling problems, while the activity in the same regions during monetary reward anticipation showed a moderate (r=-0.37) negative association with duration of abstinence. All other correlations were relatively low.

#### Relation between striatal ‘ups’ and ‘downs’

No significant (negative) correlations were found between striatal activity during cue-reactivity (hypothesized ‘up’) and monetary reward anticipation (hypothesized ‘down’) in either AUD or GD patients (Figure 4A). Bayes Factors provided anecdotal evidence in AUD patients and substantial evidence in GD patients for an absence of such a relationship (Supplementary Tables 3 and 4), indicating that these factors are independently present across individual patients.

**Figure 4.**
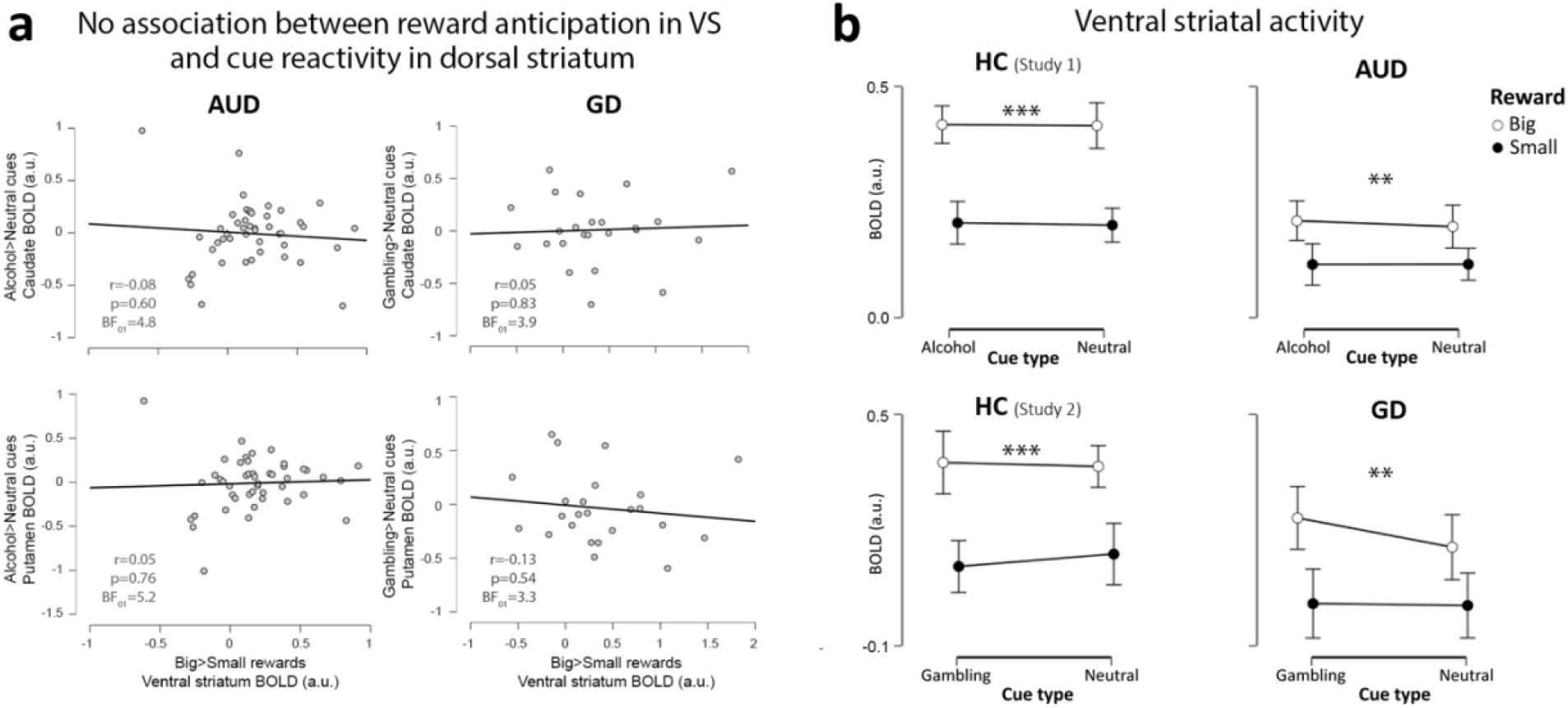
Striatal region of interest analyses, showing the (absence of a) relationship between striatal hypo- and hyperactivity in the AUD and GD group (a), and ventral striatal activity for the four experimental conditions (b). **(a)** No correlation was found between hypoactivity (big>small) and hyperactivity (addiction-related>neutral cues) in either the AUD group or the GD group. BF_01_ reflects Bayes Factors in favor of the null, reflecting how much more likely these data are to be observed under the hypothesis that there is no relationship between striatal hyperactivity and hypoactivity. **(b)** Ventral striatal activity during the four conditions, plotted separately for the patient groups and matched controls. Activity was significantly higher for big than small rewards in all groups. AUD patients showed significantly decreased activity during the processing of big rewards compared to HCs, i.e. hypoactivity. \*\*\**p*<0.001, \*\**p*<0.01, BOLD = blood oxygenation level dependent, a.u = arbitrary units.

## DISCUSSION

The present study assessed monetary reward anticipation in the presence versus absence of addiction-related cues in AUD and GD patients compared to HCs. Our results suggest that, compared to HCs, currently abstinent participants with AUD exhibited diminished striatal responses during monetary reward anticipation. Moreover, AUD patients showed increased neural responses to alcohol cues, but these activation patterns did not significantly differ from HCs. Participants with GD did not show decreased striatal neural responses during the anticipation of monetary rewards or increased activity in response to addiction-related cues compared to HCs.

In line with our hypothesis and previous studies (Beck et al., 2009; Wrase et al., 2007), AUD showed significantly decreased striatal responses during monetary reward anticipation compared to HCs. However, relative to HCs, participants with GD only had decreased activity in temporal regions but no differences in striatal activity. Following previous findings in both AUD (Chase, Eickhoff, Laird, & Hogarth, 2011; Schacht, Anton, & Myrick, 2013; Sjoerds et al., 2014) and GD (Goudriaan et al., 2010; Limbrick-Oldfield et al., 2017), both groups showed increased activity in the presence of addiction-related compared to neutral cues in a number of regions previously implicated in cue-reactivity (Schacht et al., 2013), including the ACC, precuneus (Courtney, Ghahremani, London, & Ray, 2014), insula (Limbrick-Oldfield et al., 2017) and visual areas (Hanlon, Dowdle, Naselaris, Canterberry, & Cortese, 2014). However, these neural patterns were not significantly different from the responses observed in HCs. There are several potential explanations for the unexpected lack of cue-elicited striatal activation. First, our findings are in line with the results from a meta-analysis of alcohol cue-reactivity studies, which found that striatal activity in response to alcohol cues does not differentiate cases from controls (Schacht et al., 2013). Suggested reasons for this are (among others) drug availability and treatment status (Jasinska, Stein, Kaiser, Naumer, & Yalachkov, 2014). Second, the monetary rewards in our design may have interfered with the cue-processing and thereby abolished the cue-reactivity effect. This interference could have occurred in two ways. On the one hand, the fact that the addiction cues were shown on the background could have led to attentional distraction (see limitation section for more discussion). On the other hand, money itself may already have addiction-like incentives properties: it is very closely related to the disorder in gambling, but in alcohol use disorder too, as it is fundamentally tied to the purchase of alcohol and thus to engagement in alcohol use. Thus, the appearance of the coins may have led to cue-reactivity even in the neutral condition. In sum, our findings are in line with previous work suggesting that striatal cue-reactivity is not as robust as might be assumed, and our specific task-design may have further abolished an already difficult to induce effect. A promising way forward may be to use virtual reality technology, which researchers have recently started to exploit to create naturalistic settings in cue-reactivity studies(Bruder, Scharer, & Peters, 2021; Ghiţă & Gutiérrez-Maldonado, 2018).

Behaviorally, an interaction was seen in GD patients and HCs between gambling-related cues and rewards, such that gambling cues amplified the effect of reward magnitude in both directions. Thus, participants performed better (lowest reaction time, highest accuracy) in the presence of gambling cues when playing for big rewards, but performed worse when playing for small rewards. One explanation could be that gambling cues boost participants’ impulsivity specifically when playing for bigger rewards (Miedl, Büchel, & Peters, 2014). Alternatively, this may be interpreted as an increased motivational effect of gambling cues on performance (Genauck et al., 2019), an effect also known as Pavlovian-instrumental transfer (Dickinson & Balleine, 1994), which is often quoted as an important factor for the development of and relapse to addictive behavior in associative-learning models of addiction (Everitt & Robbins, 2015; Hogarth, Balleine, Corbit, & Killcross, 2013). However, no interaction between cue-reactivity and reward anticipation or any relation with craving was seen on a neural level.

Abnormal striatal processing may constitute a biomarker of addiction severity and risk for relapse (Courtney, Schacht, Hutchison, Roche, & Ray, 2016; Jasinska et al., 2014). Several studies have previously reported correlations between various addiction-related factors and striatal activity during MIDT (reviewed in Balodis & Potenza, 2015) or cue-reactivity tasks (reviewed in Schacht et al., 2013; Starcke et al., 2018), although often using small (n<15) samples. In our exploratory analyses, we tested these relations and despite our relatively large AUD sample, we did not replicate associations between striatal activity and addiction severity (Sjoerds et al., 2014), abstinence duration (Li et al., 2014) or relapse (Courtney et al., 2016). Notably, however, AUDs who reported having more obsessive alcohol-related thoughts (higher score on OCDS) did show lower reward anticipation levels in several areas of the reward circuitry including the dorsal striatum, replicating previous findings (Wrase et al., 2007).

To the best of our knowledge, this is the first study to directly investigate the relationship between striatal ‘ups’ (related to addiction cues) and ‘downs’ (related to monetary reward anticipation). No significant relation was found between ‘striatal ups and downs’, and this independence may be interpreted to suggest that they are no related, independent factors. However, this absence should be interpreted with caution, given that we did not see robust striatal cue-reactivity effects on a group-level, and thus the operationalization of cue-reactivity within the MIDT task was limited. Interestingly, a recent study found that measures of subjective alcohol reward are related to neural indices of monetary reward in non-addicted humans (Radoman et al., 2021). They found that participants who reported greater motivation (i.e., wanting) to consume more alcohol after a single moderate dose of alcohol also exhibited greater neural activation in the bilateral ventral caudate and the nucleus accumbens during reward receipt relative to loss, on a subsequent fMRI session (Radoman et al., 2021). Hence, in non-addicted populations, there might be a relation between striatal sensitivity to drug rewards and striatal monetary rewards, which may get dissociated during the development of an addiction.

Several limitations need to be considered when interpreting these results. First, our patients were all abstinent patients recently treated with cognitive behavioral therapy in which techniques against craving are trained (e.g., craving surfing), which might have accounted for the unexpected absence of cue-elicited striatal activation in our clinical group, as suggested by a recent voxel-wise fMRI meta-analysis (Zeng et al., 2021). Second, the number of smokers was significantly higher in the AUD group than in the matched control group, while participants in the GD group differed from the HCs on IQ and handedness, which may have impacted the effects. Third, our fMRI task was designed to test the simultaneous processing of addiction-related cues and monetary rewards, but this might be problematic for the GD group as monetary rewards are the very outcomes reinforcing gambling and money may become a conditioned ‘cue’ itself. This difference between GD and AUD was one of the reasons we did not directly do comparisons between the clinical groups but rather have two separate experiments. The task-context in which monetary cues and reward anticipation are presented, may have largely differential effects in GD (van Holst, Veltman, Büchel, van den Brink, & Goudriaan, 2012; Wagner, Mathar, & Peters, 2021), but also in AUD: for instance, in an earlier studies from our group, reward expectation during a gambling task induced higher activity in the striatum in AUD compared to healthy controls (van Holst, Clark, Veltman, van den Brink, & Goudriaan, 2014). Future research could combine cue-reactivity paradigms with natural rewards (e.g. erotic cues (Sescousse, Barbalat, Domenech, & Dreher, 2013)) to investigate the imbalance of natural reward sensitivity.

In addition to money being a complex cue, a potential caveat of our task is that motivational attention is also associated with the stratal circuitry irrespective of the presence of a current or prospective reward (Boehler et al., 2011; Breckel, Giessing, & Thiel, 2011; Fan, Hof, Guise, Fossella, & Posner, 2008). Hence, reduced striatal recruitment by reward anticipatory cues in our task could actually stem from inattention to the cues rather than any intrinsic ambivalence to rewards. That said, the behavioral effect of monetary rewards on reaction time (faster responses to bigger vs smaller monetary rewards), refutes this idea. The addiction cues in our task were shown as task-irrelevant background distractors and may have captured attention away from an actual reward elicited instrumental behavior signal or vice versa, reducing the impact of the addiction cue to elicit cue-reactivity, which seems more likely given our results. Moreover, previous studies have found that drug cues can evoke activations evens when participants are unaware of having seen the cues (Childress et al., 2008). For the above-mentioned reasons and the fact that drug availability is important to generate cue-reactivity, future studies aiming at understanding the influence of addiction cues on reward anticipation could benefit from including cues for high or low monetary outcomes (so abstract cues) and also including high or low addiction relevant stimuli (such as high/low alcohol amounts), that would actually be given directly after the task (similar as the monetary outcome).

In conclusion, this first attempt to directly investigate striatal ups and downs in addictive disorders revealed that patients in treatment for AUD or GD show only some striatal hypoactivations during reward anticipation in a MIDT task, but no hyperactivation during cue-reactivity. Moreover, these factors were present independently across individuals with addiction. The finding that decreased activity during reward anticipation was seen in alcohol use but not gambling disorder is in line with the idea that repeated drug use leads to reward deficiency, while we did not find evidence for a sensitized ‘hyperdopaminergic’ striatal response to addiction cues.

## Acknowledgements

TvT, RJvH and AEG conceived and designed the study. TvT acquired, analyzed and interpreted the data, with support of RJvH and AEG. TvT prepared the manuscript; all other authors provided critical revision of the manuscript for important intellectual content. All authors read, corrected and approved the final manuscript.

This work is supported by a NWO-ZonMw grant VIDI [grant number 91713354] to A.E.G. The authors thank prof. Wim van den Brink for insightful comments on this manuscript.

