## Supplemental information for "Striatal ups or downs? Neural correlates of monetary reward anticipation, cue reactivity and their interaction in alcohol use disorder and gambling disorder"

### Supplementary Methods

#### <u>Study 1</u>

##### Participants

Initially, a total of 51 recently detoxified patients with AUD and 32 matched controls were recruited for this study. Five participants (three AUD patients and two controls) were excluded after participation because of excessive motion (>3.5mm/degrees movement from reference scan), and another two AUD patients due to scanner failure.

The relatively large AUD group was included to study the association between striatal dysfunction and individual variation in AUD severity, lifetime alcohol use, craving, abstinence and relapse. Our study was originally designed to include and compare a non-chronic (<2 years AUD history; <2 treatments) and chronic (>7 years AUD history; >3 treatments) AUD group. However, recruiting patients meeting these exact (arbitrary) cut-offs for AUD history and number of treatments was untenable. Therefore, the AUD criteria were changed during the study to include a broad range in years of AUD history, number of treatments and lifetime alcohol intake. This facilitated the investigation of chronicity by looking at severity and duration of AUD within the patient group.

**Exclusion criteria**: being younger than 18 or older than 65 years; lifetime history of bipolar disorder, anxiety disorder, obsessive-compulsive disorder or schizophrenia; past 6-month history of major depressive episode; current or past-year substance use disorder or current psychiatric treatment (except for AUD in the AUD patient group); the use of any psychotropic medication; positive breathalyzer test (alcohol) or urinalysis (benzodiazepines, (meth)amphetamines, opioids, cocaine, ecstasy, PCP, methadone or cannabis); history or current treatment for neurological disorders; major physical disorders; history of brain trauma; or any contraindications for MRI. Controls were excluded if they scored higher than 7 on the Dutch Alcohol Use Disorder Identification Test (AUDIT: Saunders, Aasland, Babor, de la Fuente, & Grant, 1993).

One AUD patient had a positive urine screen for MDMA; the subject declared that use was >1 week ago and he did not use regularly. The absence of (a history of) dependence was further verified by the psychiatric interview, and the subject was included in the analyses.

**Clinical and Neuropsychological Testing:** Verbal IQ was estimated with the Dutch Adult Reading Test (DART: Schmand, Bakker, Saan, & Louman, 1991). The WAIS digit-span (Wechsler, 1981) was used to assess general information processing speed. AUD severity was assessed with the AUDIT [score range of 0–40] (Saunders et al., 1993) and alcohol craving severity by the five-item Obsessive-Compulsive Drinking Scale [score range of 0–20] (OCDS: De Wildt et al., 2005). Lifetime alcohol intake was estimated by multiplying the total number of drinks, obtained using the Lifetime Drinking History questionnaire (Skinner & Sheu, 1982), with 0.014 (kg pure alcohol in a standard drink). Six months following participation, AUD patients were contacted by telephone to assess relapse rates, defined as >2 days of uncontrolled drinking/gambling.

Participants were asked to abstain from eating for three hours prior to the start of the assessments, because of another fMRI task (van Timmeren et al., 2020) that involved earning snack foods.

**Supplementary Table 1.**

| | **AUD (n=46)** *Mean (SD)* | **HC (n=30)** *Mean (SD)* | *p value* | *BF₁₀* |
|---|---|---|---|---|
| Age, years | 46.5 (10.8) | 44.8 (9.9) | 0.48 | 0.30 |
| Handedness: right / left | 35 / 11 | 26 / 4 | 0.26[a] | 0.58[c] |
| Males / females | 36 / 10 | 25 / 5 | 0.59[a] | 0.37[c] |
| Education, years | 10.6 (3.3) | 9.8 (3.5) | 0.21 | 0.48 |
| IQ | 102.5 (15.0) | 97.0 (9.6) | 0.07[b] | 0.87[d] |
| Smokers (%) | 30 (65%) | 4 (13%) | **<0.001**[a] | **>100**[c] |
| AUDIT | 23.7 (6.0) | 4.1 (2.1) | **<0.001**[b] | **>100**[d] |
| Lifetime alcohol intake, kg pure alcohol | 594.8 (495.8) | 52.3 (52.2) | **<0.001**[b] | **>100**[d] |
| OCDS | 5.0 (2.9) | 0.9 (1.8) | **<0.001** | **>100** |
| AUD duration (years) | 12.0 (11.0) | - | - | - |
| Weeks abstinent | 8.2 (7.6) | - | - | - |

**Supplementary Table 1:** Demographical & Clinical information AUD patients and matched controls. AUD, Alcohol Use Disorder group; HC, Healthy Control group; SD, Standard Deviation; IQ, Intelligence Quotient; AUDIT, Alcohol Use Disorders Identification Test; OCDS, Obsessive Compulsive Drinking Scale; [a]Chi-square test. [b]Mann-Whitney U test; [c]Bayesian contingency table using joint multinomial test; [d]Bayesian Mann-Whitney U (1000 samples). $BF_{10}$: Bayes Factor that quantifies the relative evidence in favor of the null (i.e. groups do not differ, $BF_{10}<1$) or the alternative (groups differ, $BF_{10}>1$) hypothesis, with $BF_{10}$ between 1 and 3 reflecting anecdotal evidence and higher than 100 extreme evidence.

#### Study 2

##### Participants

We included 24 GD patients and 22 controls in our analyses; data were collected for 26 GD patients and 23 controls, but three (two GD patients and one control) were excluded because of excessive motion (>3.5mm/degrees movement from reference scan). Patients with GD were recruited from a local addiction treatment centre (Jellinek, Amsterdam) and included if they were diagnosed with and started therapy for GD. All subjects underwent the MINI structured psychiatric interview (Sheehan, Lecrubier, & Sheehan, 1998), to confirm the absence of psychiatric disorders (except for DSM-5 GD in the GD patient group). Inclusion and exclusion criteria were furthermore identical to Study 1. One patient tested positive on THC use but was included for further analysis because drug use occurred >72 hours before inclusion and the absence of (a history of) marihuana dependence was further verified by the psychiatric interview.

**Clinical and Neuropsychological Testing:** The past-12-month Problem Gambling Severity Index (PGSI; Ferris and Wynne, 2001) and the Gamblers' Beliefs Questionnaire (GBQ; Steenbergh et al., 2002) were used to assess GD severity in GD patients. The GBQ contains 21 items (e.g. "My choices or actions affect the game on which I am betting" or "I am pretty accurate at predicting when a 'win' will occur"), with higher scores reflecting more gambling related distortions.

These data were collected as part of a larger two-day study protocol, including stress manipulation on a separate testing day. The fMRI task was preceded by questionnaires, neuropsychological testing and a reward-related decision making fMRI task and followed by a resting-state fMRI scan (reported in van Timmeren, Zhutovsky, van Holst, & Goudriaan, 2018). Subjects were reimbursed with 100 euros plus additional task earnings. All data was collected between May 2016 and August 2017. The scanner session started in the afternoon between 1:50 and 7:20 pm.

**Supplementary Table 2.**

| | GD (n=24)<br>*Mean (SD)* | HC (n=22)<br>*Mean (SD)* | *p value* | $BF_{10}$ |
|---|---|---|---|---|
| Age, years | 35.5 (12.4) | 35.4 (15.8) | 0.99 | 0.29 |
| Handedness: right / left | 22 / 2 | 15 / 7 | **0.01**[a] | **>100**[b] |
| Males / females | 19 / 5 | 13 / 9 | 0.14[a] | 1.36[b] |
| Education, years | 8.1 (3.1) | 9.6 (5.0) | 0.22 | 0.55 |
| IQ | 88.6 (8.9) | 96.6 (9.6) | **0.01** | **7.62** |
| Smokers (%) | 11 (46%) | 5 (23%) | 0.10[a] | 1.81[b] |
| AUDIT | 4.6 (4.3) | 3.0 (2.3) | 0.12 | 0.78 |
| PGSI (12 months) | 15.1 (5.1) | 0.0 (0.0) | **<0.001** | **>100** |
| GBQ | 86.6 (23.9) | 38.2 (19.7) | **<0.001** | **>100** |
| GD duration (years) | 10.7 (9.6) | - | - | |
| Weeks abstinent | 17.5 (22.3) | - | - | |

**Supplementary Table 2:** Demographical & Clinical information GD patients and matched controls. GD, Gambling Disordered patients; HC, Healthy Controls; SD, Standard Deviation; IQ, Intelligence Quotient; AUDIT, Alcohol Use Disorders Identification Test; PGSI, Problem Gambling Severity Index; GBQ, Gamblers' Beliefs Questionnaire; [a]p value of chi-square test; [b]Bayesian contingency table using joint multinomial test.. $BF_{10}$: Bayes Factor that quantifies the relative evidence in favor of the null (i.e. groups do not differ, $BF_{10}<1$) or the alternative (groups differ, $BF_{10}>1$) hypothesis, with $BF_{10}$ between 1 and 3 reflecting anecdotal evidence and higher than 100 extreme evidence.

#### Task instructions and procedure

Participants were instructed to respond to a target as quickly as possible in order to gain monetary rewards. They were trained until criterion (three correct responses) before entering the scanner to assure a basic understanding of the task. Figure 2 shows a schematic overview of the experimental paradigm. During each trial, participants could earn 1 or 50 cents, indicated by a coin overlaid on a background picture (alcohol or neutral). Participants were instructed to pay special attention to the background pictures, as they would have to answer questions about them after the experiment. Next, a crosshair was shown and participants were instructed to respond as fast as possible to the target. Target duration was adjusted through an adaptive algorithm (-10ms following hit, +20ms following miss) for each condition individually, such that participants should succeed on ~66% of the trials. Premature and late responses were defined as misses. Feedback about current and cumulative earnings was provided, followed by a fixation cross before a new trial started.

The task was programmed in E-Prime 2.0 software (Psychology Software Tools, Pittsburgh, Pennsylvania) and presented in the scanner on a 32 LCD BOLDscreen (Cambridge Research Systems). Participants were told they should use their right index finger to respond on the MRI-compatible button box (CurrentDesigns).

**MRI acquisition**

Magnetic resonance imaging (MRI) was performed on a 3 Tesla, full-body Intera MRI scanner (Philips Medical Systems, Best, The Netherlands) equipped with a 32-channel phased array SENSE radiofrequency (RF) receiver head coil. A high-resolution T1-weighted structural image was acquired for each participant (6.862 ms repetition time; 3.14 ms echo time; 8º flip angle; 1x1x1 mm voxel size; 236.679 x 180 x 256mm field of view; 212 x 212 matrix size; 150 slices; 1.2 mm slice thickness). Functional MRI scans were acquired using a T2*-weighted gradient multi-echo echoplanar imaging sequence (2375 ms repetition time; 9 / 26.4 / 43.8 ms echo times; 76º flip angle; 3x2.95x3 mm voxel size; 76 x 73 matrix size; 37 slices, acquired in interleaved order; 3mm slice thickness; 0.3mm slice gap). This sequence was chosen for its improved blood oxygen level dependent (BOLD) sensitivity and lower susceptibility for artefacts, especially for ventral regions (Poser, Versluis, Hoogduin, & Norris, 2006). The first three scans were discarded to allow T1 saturation to reach equilibrium.

**fMRI preprocessing**

Imaging data were preprocessed using SPM12 (Wellcome Trust Centre for Neuroimaging, London). Raw multi-echo data were combined as reported in van Timmeren et al. (van Timmeren, Zhutovsky, van Holst, & Goudriaan, 2018). In short, realignment parameters were estimated for the images acquired at the first echo time and consequently applied to images resulting from the two other echoes. Thirty volumes, acquired from an independent run during which a fixation cross was shown, were used to calculate the optimal weighting of echo times for each voxel by applying a PAID-weight algorithm (Poser et al., 2006). The multi-echo fMRI data were then combined into single volumes using these weightings. All functional images were subsequently slice-time corrected and co-registered with the high-resolution T1-weighted image using normalized mutual information. The high-resolution structural scan was segmented and used to normalize the slice-time corrected functional images. Finally, all functional images were smoothed with an 8mm isotropic full-width at half maximum (FWHM) Gaussian smoothing kernel.

### Supplemental results

#### ROI analyses

**Study 1: Regional ventral striatal BOLD – AUD:** A repeated measures ANOVA indicated higher activity for big than small rewards over the groups (main effect of reward: $F_{1,74}$=78.6, p<0.001), a reward X group interaction ($F_{1,74}$=13.9, p<0.001), driven by lower activity for big rewards in the AUD group compared to the HCs ($t_{74}$=-2.96, p=0.004); and generally lower activity in the AUD patients than in the HCs (main effect of group: $F_{1,74}$=48.4, p=0.028).

**Study 2: Regional ventral striatal BOLD – GD:** Over both groups, ventral striatal activity was higher for big than small rewards (main effect of reward: $F_{1,74}$=78.6, p<0.001); however, no significant effects of group (lowest: 0.09) or cue-type (lowest: 0.08) were found.

**Sup Table 3: Correlation between striatal hyperactivity (related to addiction cues) and hypoactivity (related to monetary reward anticipation) in AUD individuals.** $BF_{01}$ reflect Bayes Factors in favor of the null, reflecting how much more likely these data are to be observed under the hypothesis that there is no relationship between striatal hyperactivity and hypoactivity. VS= Ventral striatum, Alc=alcohol cues, neu=neutral cues, big=big reward (50 cents), small=small rewards (1 cent).

| | ***VS (alc>neu)*** | | | ***Caudate (alc>neu)*** | | | ***Putamen (alc>neu)*** | | |
|---|---|---|---|---|---|---|---|---|---|
| | r | *p* | $BF_{01}$ | r | *p* | $BF_{01}$ | r | *p* | $BF_{01}$ |
| ***VS (big>small)*** | -0.22 | 0.15 | 2.0 | -0.08 | 0.60 | 4.8 | 0.05 | 0.76 | 5.2 |
| ***Caudate (big>small)*** | -0.24 | 0.12 | 1.7 | -0.22 | 0.14 | 1.9 | -0.11 | 0.49 | 4.3 |
| ***Putamen (big>small)*** | 0.04 | 0.80 | 5.3 | 0.06 | 0.68 | 5.0 | 0.19 | 0.20 | 2.5 |

**Sup Table 4: Correlation between striatal hyperactivity (related to addiction cues) and hypoactivity (related to monetary reward anticipation) in GD individuals.** $BF_{01}$ reflect Bayes Factors in favor of the null, reflecting how much more likely these data are to be observed under the hypothesis that striatal hyperactivity and hypoactivity are not related. VS= Ventral striatum, Alc=alcohol cues, neu=neutral cues, big=big reward (50 cents), small=small rewards (1 cent).

| | ***VS (alc>neu)*** | | | ***Caudate (alc>neu)*** | | | ***Putamen (alc>neu)*** | | |
|---|---|---|---|---|---|---|---|---|---|
| | r | *p* | $BF_{01}$ | r | *p* | $BF_{01}$ | r | *p* | $BF_{01}$ |
| ***VS (big>small)*** | -0.06 | 0.78 | 3.8 | 0.05 | 0.83 | 3.9 | -0.13 | 0.54 | 3.3 |
| ***Caudate (big>small)*** | -0.11 | 0.60 | 3.5 | -0.16 | 0.45 | 3.0 | -0.24 | 0.27 | 2.2 |
| ***Putamen (big>small)*** | -0.04 | 0.86 | 3.9 | -0.23 | 0.29 | 2.3 | -0.18 | 0.40 | 2.8 |

**Suppl table 5**: Correlation between various clinical measures and neural activity during Monetary reward anticipation (big > small) in the three striatal ROIs in AUD (study 1). Values represent Pearson correlation coefficients (r). The strength and direction are color coded.

-1.0 0. 1.0

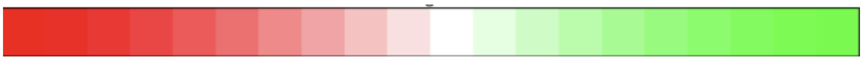

**AUD: Monetary reward anticipation (big > small)**

| | Ventral Striatum | Caudate | Putamen |
|---|---|---|---|
| Alcohol craving | -0,14 | -0,06 | -0,15 |
| AUD severity | -0,06 | -0,15 | -0,20 |
| AUD duration | 0,08 | -0,01 | 0,01 |
| Abstinence | 0,04 | 0,17 | -0,02 |
| OCDS | -0,24 | -0,27 | -0,44 |
| Alcohol intake (kg) | 0,07 | 0,15 | 0,16 |

**Suppl table 6**: Correlation between various clinical measures and neural activity during cue-reactivity (alcohol-related > neutral cue) for the three striatal ROIs in AUD (study 1). Values represent Pearson correlation coefficients (r).

**AUD: Cue-reactivity (alcohol-related > neutral cue)**

| | Ventral Striatum | Caudate | Putamen |
|---|---|---|---|

| | | | |
|---|---|---|---|
| Alcohol craving | 0,05 | -0,02 | 0,05 |
| AUD severity | 0,18 | 0,27 | 0,21 |
| AUD duration | 0,08 | 0,03 | 0,05 |
| Abstinence | -0,07 | -0,02 | 0,02 |
| OCDS | -0,09 | 0,01 | 0,04 |
| Alcohol intake (kg) | -0,03 | -0,08 | 0,20 |

**Suppl table 7**: Correlation between various clinical measures and neural activity during Monetary reward anticipation (big > small) in the three striatal ROIs in GD (study 2). Values represent Pearson correlation coefficients (r).

| | **PG: Monetary reward anticipation (big > small)** | | |
|---|---|---|---|
| | Ventral Striatum | Caudate | Putamen |
| Gambling craving | -0,22 | -0,11 | -0,15 |
| GD severity | -0,06 | -0,22 | -0,25 |
| GD duration | -0,21 | -0,03 | 0,13 |
| Abstinence | -0,37 | -0,23 | -0,18 |

**Suppl table 8**: Correlation between various clinical measures and neural activity during cue-reactivity (alcohol-related > neutral cue) for the three striatal ROIs in GD (study 2). Values represent Pearson correlation coefficients (r).

| | **PG: Cue-reactivity (gambling-related > neutral cue)** | | |
|---|---|---|---|
| | Ventral Striatum | Caudate | Putamen |
| Gambling craving | -0,11 | 0,04 | -0,07 |
| GD severity | 0,25 | 0,13 | 0,00 |
| GD duration | 0,44 | 0,06 | 0,08 |
| Abstinence | 0,14 | 0,21 | 0,17 |

**Figure S1**: Accuracy (number of correct trials) in (A) AUDs + matched HCs and (B) GDs + matched HCs

AUD group

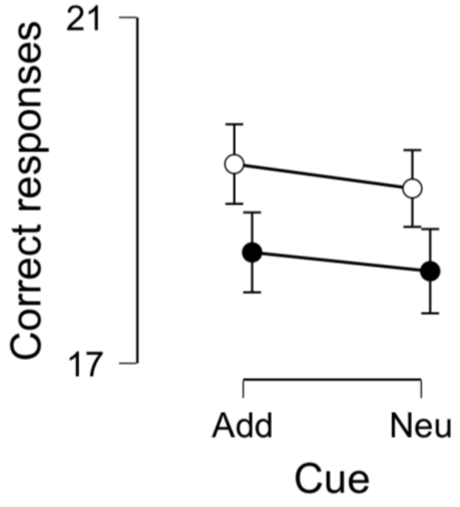


HC group (AUD study)

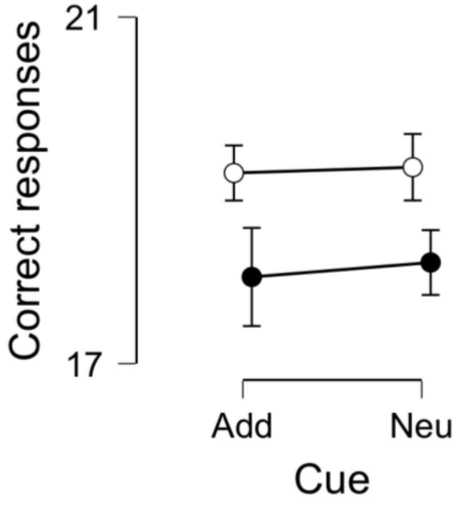


GD group

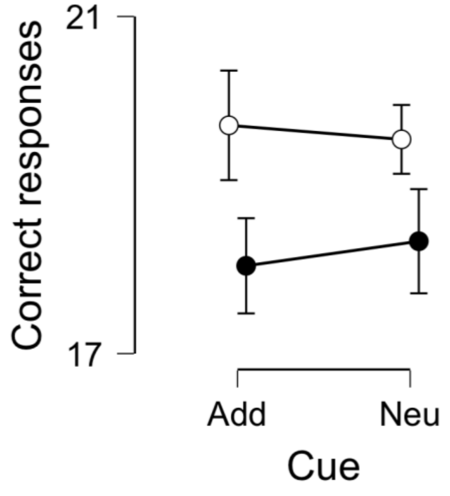


HC group (GD study)

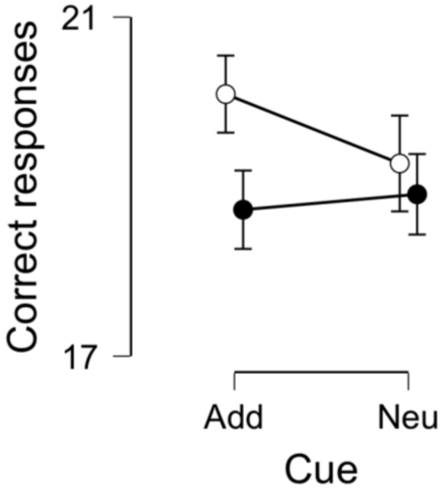

**Figure S2:** Whole-brain statistical parametric map for the cue reactivity effect within the GD group.

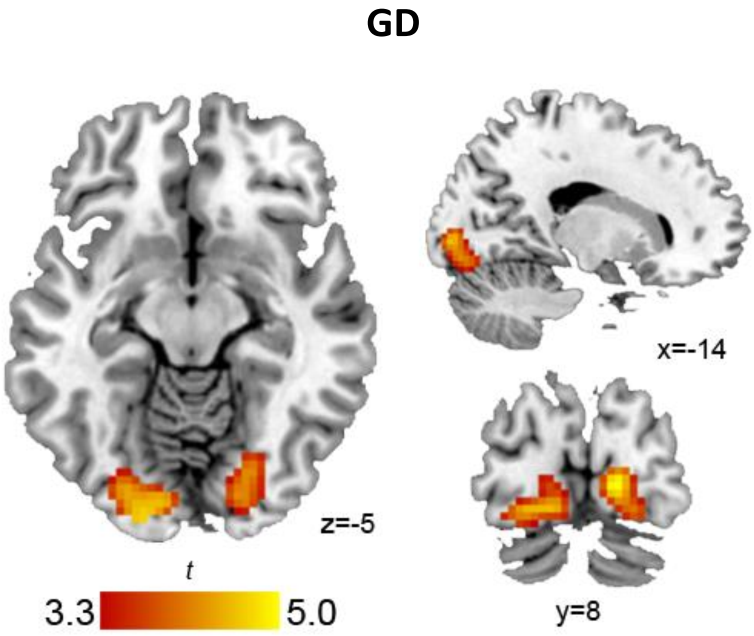

**Figure S3. Results from the caudate and putamen regional ROI analyses:** activity (betas, arbitrary units) on the y-axis during the four conditions, plotted separately for each group for the caudate and putamen.

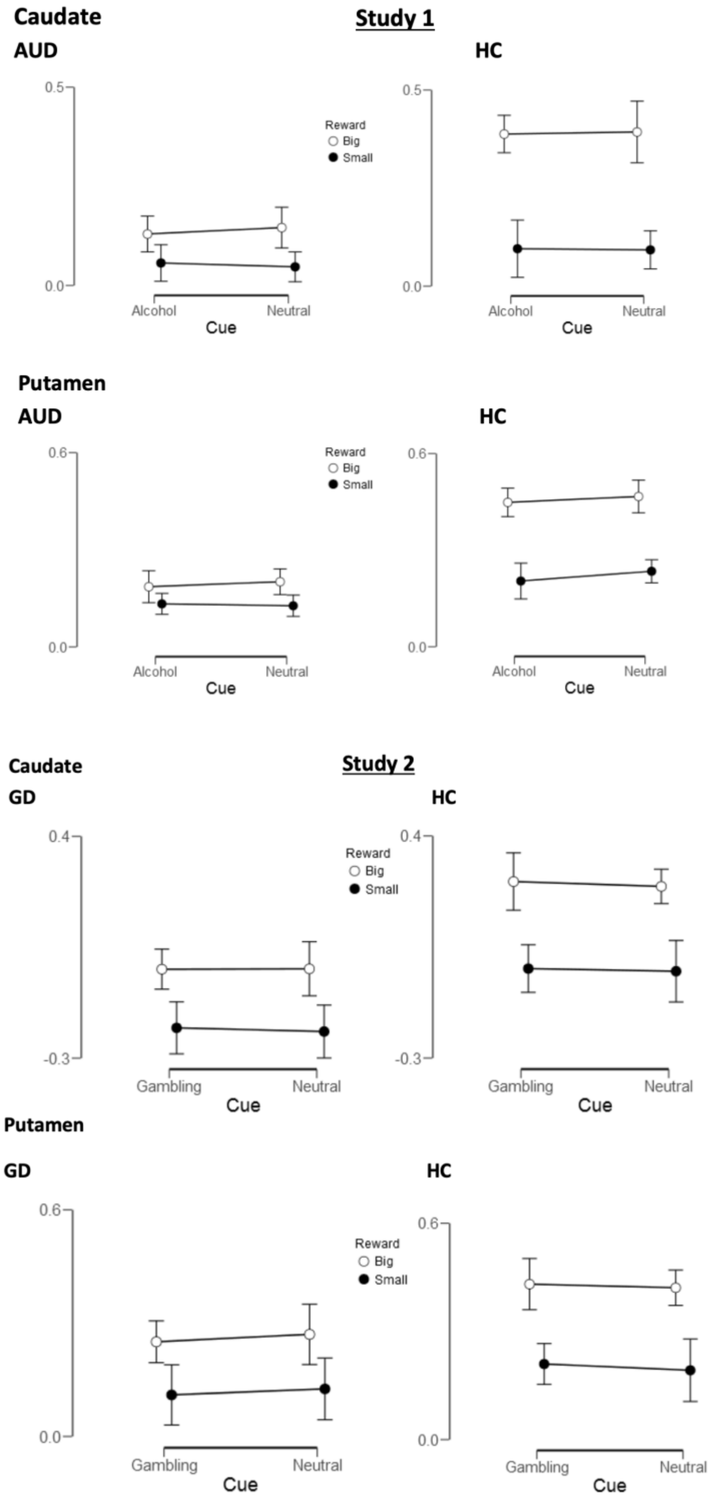
